# Task-dependent modulation of spinal and transcortical stretch reflexes linked to motor learning rate

**DOI:** 10.1101/223842

**Authors:** Michael Dimitriou

## Abstract

It is generally believed that task-dependent control of body configuration (‘posture’) is achieved by adjusting voluntary motor activity and transcortical ‘long-latency’ reflexes. Spinal monosynaptic circuits are thought not to be engaged in such task-level control. Similarly, being in a state of motor learning has been strongly associated only with an upregulation of feedback responses at transcortical latencies and beyond. In two separate experiments, the current study examined the task-dependent modulation of stretch reflexes by perturbing the hand of human subjects while they were waiting for a ‘Go’ signal to move at the different stages of a classic kinematic learning task (visuomotor rotation). Although the subjects had to resist all haptic perturbations equally, the study leveraged that task-dependent feedback controllers may already be ’loaded’ at the movement anticipation stage. In addition to an upregulation of short- and long-latency reflex gains during early exposure to the visual distortion, I found a relative inhibition of reflex responses in the ‘washout’ stage (sensory realignment state). For more distal muscles (brachioradialis), this inhibition also extended to the monosynaptic reflex response (‘R1’). These R1 gains reflected individual motor learning performance in the visuomotor task. The results demonstrate that the system’s ‘control policy’ in visuomotor adaptation can also include inhibition of proprioceptive reflexes, and that aspects of this policy can affect monosynaptic spinal circuits. The latter finding suggests a novel form of state-related control, probably realized by independent control of fusimotor neurons, through which segmental circuits can tune to higher-level features of a sensorimotor task.

**Additional Information:** *Conflict of interest:* The author declares no conflict of interest.

*Author contributions:* M.D. devised, designed and implemented the project, analyzed the data and wrote the manuscript. The author approves the final version of the manuscript and is fully accountable for all aspects of the work. The experiments were performed at the Department of Integrative Medical Biology, Umeå University, Sweden.

*Funding:* This work was financially supported through grants awarded to M.D. by the Kempe Foundation, the local Foundation for Medical Research (“Insamlingsstiftelsen”) and the Swedish Research Council (project 2016-02237).

## Introduction

At least with respect to postural control, a widely-held belief for many years has been that higher-level aspects of a sensorimotor task (e.g., goal or context) affect voluntary motor control and transcortical motor reflex responses, whereas the gain of spinal (segmental) monosynaptic circuits varies only according to pre-perturbation muscle activity levels (e.g., Hammond, 1956; Marsden, Merton, & Morton, 1976; Pruszynski, Kurtzer, Lillicrap, & Scott, 2009; Pruszynski & Scott, 2012; Scott, 2012; Scott, Cluff, Lowrey, & Takei, 2015). Specifically, there has been some evidence that even the monosynaptic (‘short-latency’) neuromuscular response can modulate during cyclical behaviors such as gait (e.g., Akazawa, Aldridge, Steeves, & Stein, 1982; Capaday & Stein, 1986), or following the onset of muscle activity associated with a ballistic voluntary movement (but not before, see e.g., Mortimer, Webster, & Dukich, 1981) or after extensive training in primates (e.g., Wolpaw, 1982). However, to my knowledge, there has been no evidence of a flexible, systematic and context-dependent modulation of monosynaptic responses to postural perturbations across a group of human subjects. With regard to posture and goal-directed reaching as well, it appears the lack of such evidence has -justifiably-led researchers to assume that the monosynaptic stretch reflex is generally not task-dependent, and only affected by the perturbed muscle’s state. In other words, the short-latency response is generally thought to occur outside the nervous system’s ‘control policy’ framework which determines the preferred feedback response to sensory inflow (see e.g., Scott, 2016; Scott et al., 2015).

In the same vein, previous studies have shown that being in a state of motor learning (i.e., adapting to novel dynamics) increases or ‘upregulates’ the gain of feedback responses only at transcortical reflex latencies and beyond, with a similar upregulation or no effect when adapting to the sudden removal of the altered dynamic state (Ahmadi-Pajouh, Towhidkhah, & Shadmehr, 2012; Cluff & Scott, 2013; Franklin, Wolpert, & Franklin, 2012). However, recent recordings of proprioceptive afferent signals from distal (wrist) muscles during visuomotor adaptation (Dimitriou, 2016) have introduced the possibility of more flexible behavior during motor learning, that may include inhibition of rapid feedback responses (i.e., ‘reflexes’). Moreover, it is hypothesized that possible state-dependent tuning of proprioceptors in visuomotor learning will lead to task-related output at spinal monosynaptic latencies, demonstrating a higher degree of sophistication in such circuits than previously thought.

To test the above, subjects made center-out reaching movements with their right hand in the context of a classic visuomotor learning paradigm, while grasping the handle of a robotic manipulandum (Figure 1AB). On randomly interleaved probe trials within specific stages of the learning task (Figure 1C), the position of the hand was perturbed during the movement preparation stage, i.e., while waiting for a ‘Go’ signal to move after the target has been visually highlighted. The haptic perturbations were applied in order to examine the nature of stretch reflex responses at the different states of visuomotor adaptation (Figure 1DE: Experiment 2 differed from Experiment 1 in that a mechanical load was applied in probe trials). At least when adapting to novel dynamics, the nervous system is known to ‘load’ task-dependent feedback controllers already at the movement preparation stage (Ahmadi-Pajouh et al., 2012). Therefore, even if subjects had to resist all haptic perturbations in order to keep the hand immobile before the ‘Go’ signal was given, task-specific differences were still expected as a function of the ‘pre-loaded’ feedback controllers. The current study also allowed for an examination of individual differences in the relationship between reflex gain modulation (probe trials) and motor learning performance (non-probe trials).

**Figure 1.**
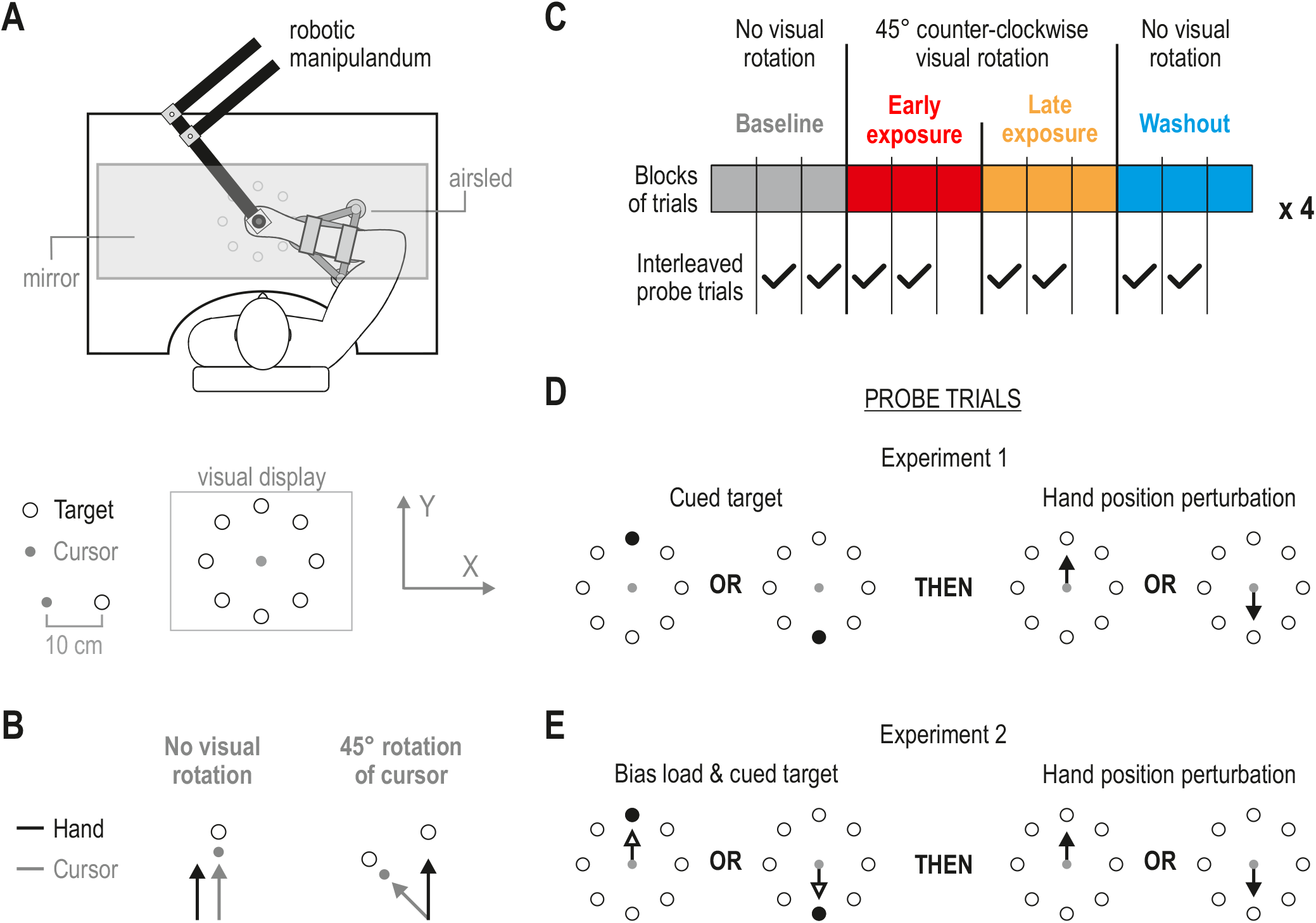
Experimental paradigm. (A) The subjects’ right forearm rested on an airsled and they grasped the handle of a robotic manipulandum. Visual feedback was projected by a monitor onto the one-way mirror that also prevented view of the right limb. The instantaneous position of the hand was represented by a small moving dot (‘cursor’). The subjects were required guide the cursor in a straight line towards visual targets by making discrete center-out movements. On each trial of this main task, a target would be visually cued (highlighted red) but subjects were instructed to remain in the central start position until a ‘Go’ signal was given (red highlighted target turned green). (B) Subjects performed the center-out movements with or without experiencing a 45° counter-clockwise rotation of the direction of the cursor. (C) Each block of trials of the main task required movement to each of the 8 possible targets (blocked-randomized presentation). The visual rotation was suddenly introduced after 3 initial blocks, remained for another 6 blocks, and was removed for the following 3 blocks (‘washout’ stage, blue). The whole process was repeated 4 times. Additional ‘probe’ trials were randomly interleaved within specific blocks (tick marks in C) in order to assess the gain (i.e., output magnitude given the same input) of stretch reflexes at the different stages of the main task. (D) In Experiment 1, the hand was unpredictably perturbed either in the same or opposite direction of a cued visual target, before the ‘Go’ signal was given (i.e., postural perturbations). Because the subjects were instructed to keep their hand inside the start point until the ‘Go’ signal, they had to resist all hand perturbations (E) Experiment 2 was the same as Experiment 1, except that a slowly-rising 6N load was applied in the direction of the subsequently cued target (probe trials only). Throughout, both the onset (timing) and direction of the haptic position perturbations were unpredictable.

## Materials and Methods

### Subjects

In total, 30 individuals took part in the current study: 15 subjects (6 males) participated in Experiment 1 (mean age of 25 ± 5 years) and 15 subjects (7 females) participated in Experiment 2 (mean age of 25.5 ± 4 years). All participants were right-handed, neurologically healthy and were financially compensated for their contribution. All gave their written consent to participating in the study prior to experimentation according to the Declaration of Helsinki. The current study was part of a program approved by the Ethics Committee of Umeå University, Umeå, Sweden.

### Experimental setup

Subjects sat in an adjustable chair with their right hand holding onto the handle of a robotic manipulandum (KINARM end-point robot, BKIN Technologies, CA). The KINARM is able to produce controlled forces on the hand, whereas the forces applied by the subject on the robotic handle are measured by a six-axis force transducer (Mini40-R, ATI Industrial Automation). The system also generates kinematic data with regard to the position of the handle. The subject’s right forearm was placed inside a thin cushioning foam structure, itself attached to a custom-made airsled, similar to that used elsewhere (Dimitriou, Wolpert, & Franklin, 2013; Howard, Ingram, & Wolpert, 2009). The airsled supported the subject’s forearm and allowed for frictionless motion in a 2D plane (Figure 1A). A piece of soft leather fabric with Velcro attachments was wrapped tightly around the forearm and hand of the subject, reinforcing the mechanical connection between the airsled, the handle and the hand. In this setup, surface Electromyography (EMG) was also recorded from six muscles of the right arm (see relevant section for more details). Visual stimuli were displayed into the plane of movement by means of a one-way mirror, on which the contents of a monitor were projected. In this standard setup the subject has no vision of their hand (Figure 1A). Instead, the instantaneous position of the subjects’ hand during voluntary movement was visually represented by a white filled circle with 1 cm diameter (‘cursor’). Visual targets were represented by circles (2 cm diameter) that were placed symmetrically at 45° intervals, each at a distance of 10 cm from a central point (Figure 1A, bottom schematic).

### Experimental paradigms

Two separate experiments were performed, but the underlying experimental approach was the same. The subjects were asked to perform a classic visuomotor learning task, with ‘probe’ trials (hand perturbation trials) randomly interleaved at different points during the main task. The main task in each experiment always required subjects to make center-out movements with their right arm, to bring the hand in a straight line to one of eight peripheral visual targets (Figure 1A). The subjects performed these discrete movements with or without experiencing a 45° counterclockwise rotation of cursor direction (Figure 1B). All movements and hand perturbations begun from a central start point (red circle, 2 cm diameter), located approximately 30 cm in front of the subject’s chest (Figure 1A). Unless otherwise indicated below, all possible visual target locations (N=8) were continuously displayed in the form of brown circular outlines (2 cm diameter). All hand position perturbations in the current study were applied in Cartesian space (+Y or -Y direction) before movement onset, and the visual position of the hand was not updated for the duration of these imposed hand displacements (i.e., cursor position was ‘frozen’). As described more detail below, all position perturbations were unpredictable to the subjects in terms of their timing (onset) and direction, disallowing any pre-programmed or volitional responses to be prepared or otherwise cued in advance. All perturbations were applied before the ‘Go’ signal was shown; that is, the perturbation occurred while the subject was still maintaining the position of the hand, as required by the task (before any muscle activity associated with transition to movement).

#### Experiment 1

Experiment 1 examined how stretch reflex gains modulate at different stages of the classic visuomotor adaptation task, by using postural perturbations of the hand to probe such responses. Specifically, subjects initiated each trial by placing the cursor -representing hand position-within the start circle located in the center of the display (Figure 1A), and waiting there immobile for 1 sec + a random time (i.e., random choice from the interval 1-500 msec). After this time, one of the eight peripheral targets was suddenly cued by turning from a brown circular outline to a red filled circle of the same diameter as the outline. The subjects were instructed beforehand to remain immobile inside the central start circle until the ‘Go’ signal was given (the red cued target suddenly changed color to green). The time between cuing a target and giving the ‘Go’ signal to move was 1 sec + a random time (from the interval 1-500 msec). Upon reaching a target, the subjects were required to leave their hand there immobile until they received visual feedback on their performance. That is, “correct” or “too slow” was shown after remaining immobile inside the target for 300 msec. The subjects received the “too slow” feedback if they took >800 msec to reach the target (calculated from the onset of the ‘Go’ signal). More detailed feedback and/or more stringent speed and accuracy requirements concerning the subject’s voluntary center-out movements was thought unnecessary for the purposes of the current study. The main reason for performing the kinematic learning task was to place the subjects in the different visuomotor adaptation states (i.e., Figure 1C). In other words, the subjects are assumed to be e.g., in a ‘baseline’ state during the ‘baseline’ stage of the task. After receiving feedback regarding their center-out movement, the subjects were then free to return to the start-point to initiate the next trial. There was no time limit associated to this return movement, so the overall task was essentially self-paced. A block of such trials required movement to one of each of the eight target locations (blocked-randomized presentation). As commonly the case, the first three blocks of trials in this process are referred as belonging to a ‘baseline’ stage (Figure 1C). Just after these first three blocks of trials were finished (i.e., on trial 25), a 45° counter-clockwise rotation of cursor direction was suddenly applied. This visual distortion remained for another 6 blocks of trials, and was then suddenly removed just before the onset of trial 73. The former three blocks (blocks 4-6) are referred to as belonging to the ‘early exposure’ stage and the latter three blocks are part of ‘late exposure’ stage. Three additional blocks of trials without the visual rotation then followed, in order to examine aftereffects (‘washout’ stage). The whole process described above (i.e., 12 blocks) was then repeated another three times (Figure 1C). In total, each subject performed 4 × 12 × 8 = 384 voluntary movement trials. The subjects were allowed to take a short break (<5 minutes) at the end of 12 consecutive blocks of trials (e.g., see Figure 2).

**Figure 2.**
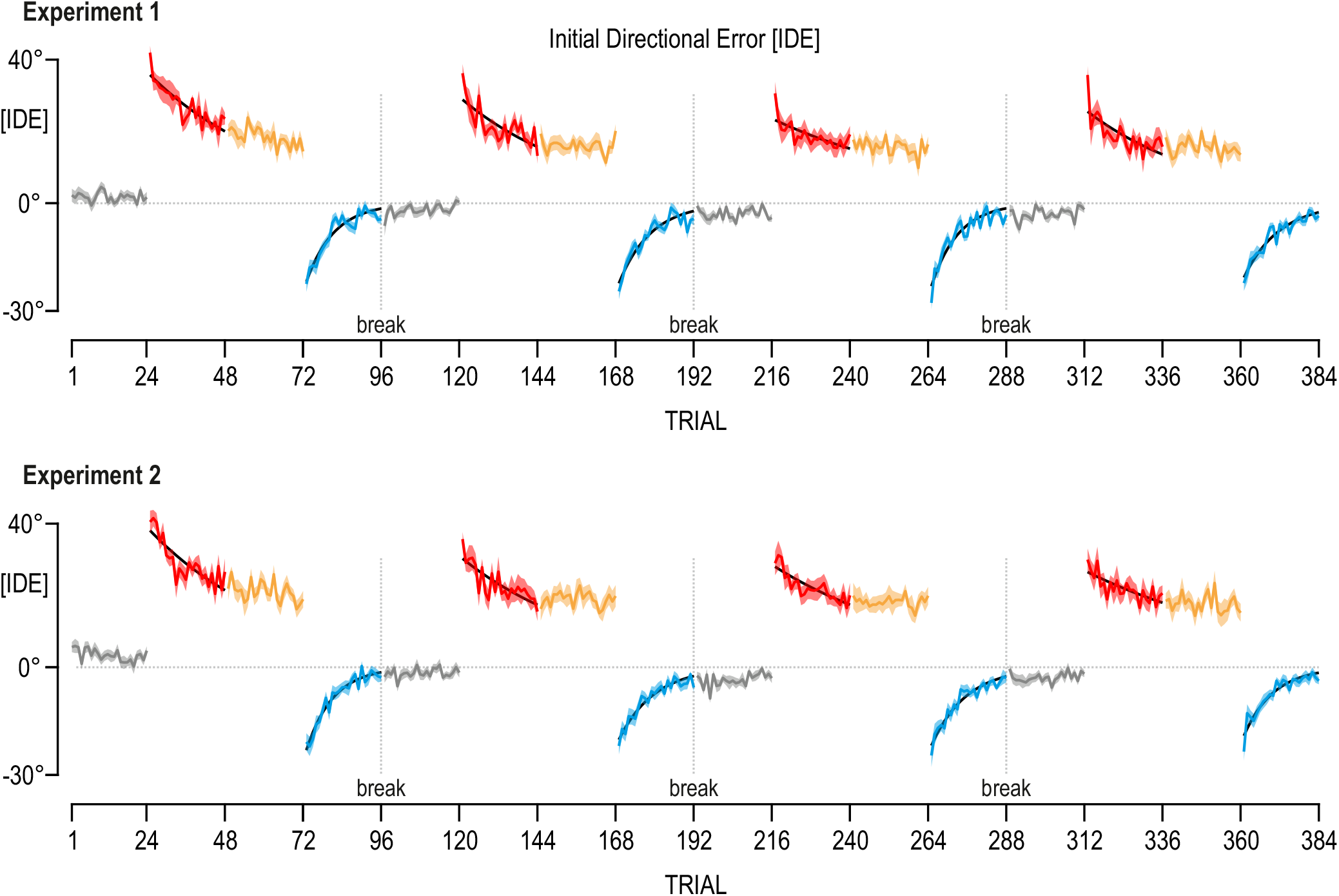
Visuomotor adaptation across subjects. To quantify adaptation rate (i.e., learning rate), Initial Directional Error (IDE) values were calculated for each non-probe trial. As commonly defined, IDE represents the angular difference between the direction the hand was moving (calculated at initial peak velocity in each movement) and the direction subjects should have been moving to, given the target location. An IDE value was therefore produced for each voluntary movement of the main visuomotor learning task in each experiment. Here, the colored traces represent the mean IDE across subjects in Experiment 1 and Experiment 2. Shaded colored areas indicate ±1 SEM. The same color scheme is used as in Figure 1. Despite repeated application and removal of the visuomotor rotation (i.e., 4 repetitions of the main process; Figure 1C), stereotypical behaviors were observed throughout. Exponential curves (thick black lines) could significantly account for progression of mean IDE in all ‘early exposure’ (red) and ‘washout’ stages (blue) (Experiment 1: all R^2^>0.55, all p<10^-5^; Experiment 2: all R^2^>0.59, all p<10^-5^). Specifically, large movement direction errors occurred in early exposure to visuomotor rotation, and such errors decayed exponentially on a trial-by-trial basis during the ‘early exposure’ stage (red). Sudden removal of the visual rotation produced IDE’s in the opposite direction in both experiments (‘after-effects’), and these errors also decayed exponentially (‘washout’ stage; blue). The results confirm that stereotypical adaptation states were induced in both experiments.

In addition to the voluntary movement trials above, probe trials were randomly interleaved within specific blocks of the visuomotor learning task: at the latter two blocks of ‘baseline’ and the initial two blocks of ‘early exposure’, ‘late exposure’ and ‘washout’ stages (tick symbols in Figure 1C). These specific blocks were chosen in order to focus on responses early in each adaptation stage (‘early exposure’ and ‘washout’), in combination with maximizing the distinguishability across the different stages. The four different types of probe trials that occurred within a block of Experiment 1 are shown in Figure 1D. As with the non-probe trials, the subjects initiated a probe trial by placing the cursor within the start circle and waiting there for 1 sec + a random time (chosen from the interval 1-500 msec). Then, either the north visual target (‘+Y’) or south target (‘-Y’) was suddenly cued (highlighted red, filled circle). After a fixed time of 950 msec + a random time (from the interval 1-300 msec), a position-controlled perturbation of the subjects hand occurred in either the +Y or -Y direction (3.5 cm displacement, 150 msec rise time and 50 msec hold). Note that the haptic perturbation occurred before the ‘Go’ signal was shown. That is, the perturbation was applied before any muscle activity associated with voluntary movement. The perturbation itself was designed to induce the kinematics of a fast naturalistic point-to-point movement (i.e., bell-shaped velocity profile), and the robot was allowed to employ maximum available stiffness (~40,000 N/m) -if necessary-to achieve the desired kinematics. The KINARM robot was able to impose the required hand kinematics of these perturbations reliably across the different stages of the main task (e.g., see Figure 3).

**Figure 3.**
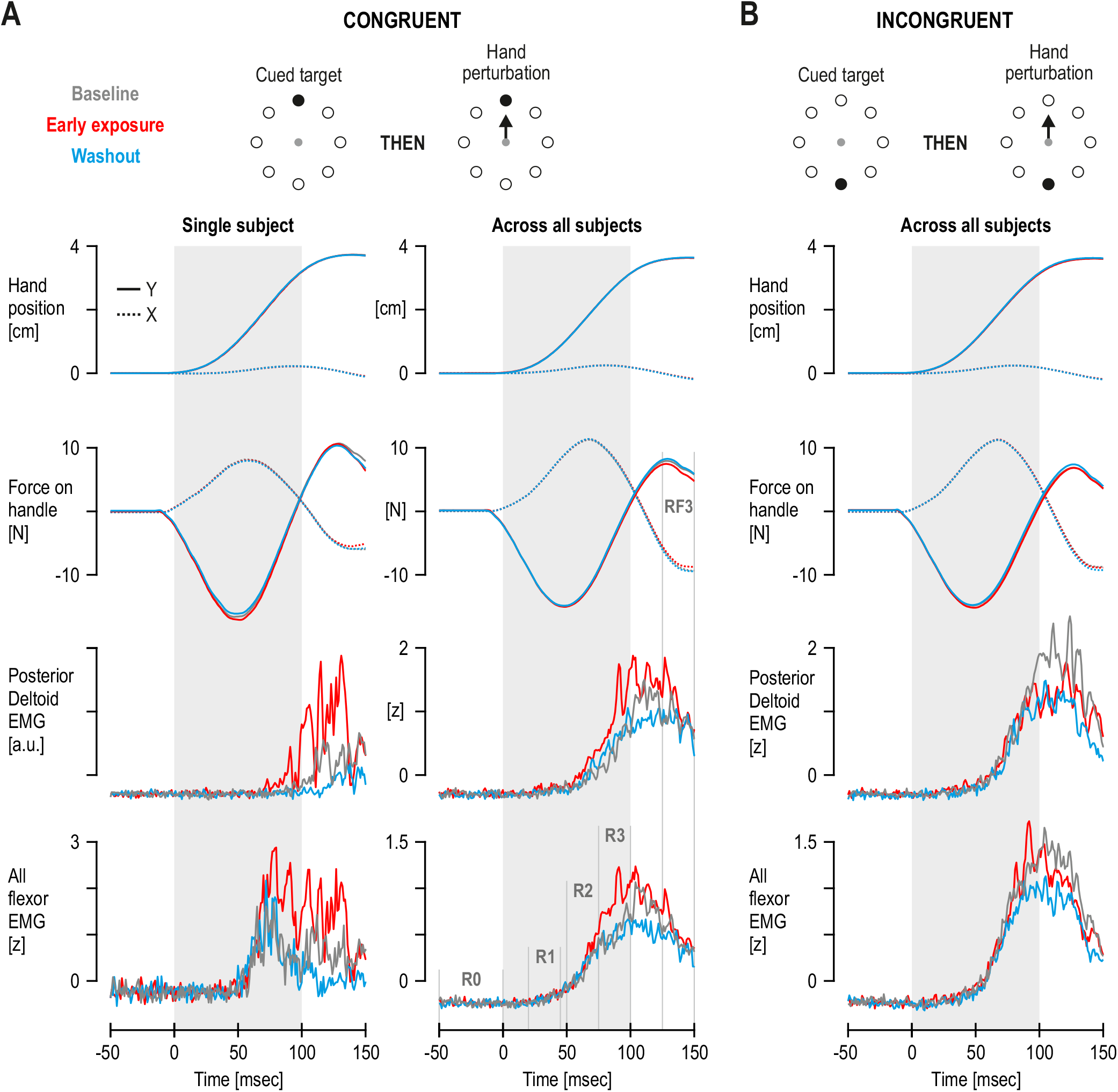
Responses to hand perturbations in Experiment 1. (**A**) Averaged single-subject responses across repetitions (left column) and corresponding mean responses across subjects (right column) when the hand was perturbed in the direction of flexor stretch (i.e., away from the torso), during different stages of the visuomotor learning task. ‘Flexor’ EMG is the average (mean) across all recorded flexor muscles of a single subject (i.e., z-transformed brachioradialis, biceps, and posterior deltoid). Here, the cued target location was congruent to the direction of the subsequent perturbation (see schematic). ‘Late exposure’ traces and error bars are omitted for visual clarity, but relevant variances across subjects are displayed in Figure 4. Despite same pre-perturbation activity levels (‘R0’) and virtually identical kinematics during all position-controlled perturbations, there are clear differences in long-latency responses as a function of task state. Specifically, both at the individual muscle level and across all recorded flexors, the ‘R3’ EMG response (75 -100 ms) during the ‘early exposure’ state (‘red’) was higher than those observed in either ‘baseline’ or ‘washout’. (**B**) As right column of A, except the cued visual target was in opposite direction of the subsequent haptic perturbation (see schematic). Here, the overall EMG responses to the same ‘+Y’ perturbation were higher across all stages of the task compared to the case where perturbation direction and cued target location were congruent. This effect essentially replicates previous findings, where a very similar manipulation was applied outside the context of visuomotor learning. However, the differences in flexor EMG across visuomotor adaptation stages within this latter case appear less clear compared to the former (i.e., ‘B’ vs. ‘A’).

Because the four probe trials were randomly interleaved among the eight non-probe trials within a block, the occurrence, timing and direction of any perturbation was unknown to the subjects, even after a target was cued. The subjects had to resist any such postural perturbation because, as mentioned above, they were instructed to remain immobile inside the central start circle until they received a ‘Go’ signal. After the perturbation ended (i.e., after position-control of the handle was removed; force ramp-down time of 20 msec), the subjects swiftly returned their hand to the start circle. After a fixed time of 0.5 sec + a random time (from the interval 1-500 msec), the ‘Go’ signal was given and the subject then moved to the target. The trial ended when the subjects kept their hand immobile inside the target until they received visual feedback (“correct” or “too slow”), as per the non-probe trials. After receiving feedback, the subjects returned to the start-point to initiate the next trial, which may or may not have been another probe trial. The total number of probe trials experienced by a single subject was 4 × 8 blocks × 4 trial types = 128 probe trials. That is, a total of 8 repetitions per probe trial type was obtained, per task stage (Figure 1CD). The subjects therefore performed 384 (non-probe trials) + 128 (probe trials) = 512 trials in total. The whole experimental session lasted ~1.5 hours.

#### Experiment 2

A second experiment examined whether the application of interleaved mechanical loads during the kinematic learning task allows for the observation of flexible task-dependent responses even at monosynaptic latencies (commonly used experimental manipulation for this purpose). Experiment 2 was the same as Experiment 1, except that mechanical loading preceded the cuing of targets in probe trials alone (Figure 1E). Specifically, each probe trial begun by placing the cursor within the start circle, and waiting there for 1 sec + a random time (i.e., random choice from the interval 1-500 msec). A mechanical load (6 N) in either the +Y direction (flexor load) or -Y direction (extensor load) was then gradually applied by the handle on the subject’s hand (a slow continuous rise-time of 2 sec). Again, the subjects countered the load and maintained the hand inside the central start circle, because they were instructed to remain immobile there until the ‘Go’ signal was shown. When the load reached 6 N and the hand was immobile inside the start circle, the corresponding +Y or -Y visual target was cued (i.e., the target located along the direction of the load; Figure 1E). After a fixed time of 950 msec + a random time (from the interval 1-300 msec), a position-controlled perturbation of the subjects hand occurred in either the +Y or -Y direction. That is, the process continued exactly as in Experiment 1, with the same position-controlled perturbation occurring before the ‘Go’ signal was given. The presentation of the trials was blocked-randomized, and therefore the direction (and timing) of the postural perturbations was unknown to the subjects, even after a target was visually cued.

### Electromyography

In all experiments, the Delsys Bagnoli (DE-2.1–Single Differential Electrodes) system was used to record surface EMG from six muscles actuating the right upper limb: brachioradialis, biceps-brachii, triceps lateralis, triceps longus, the posterior deltoid and the anterior deltoid. That is, three upper limb flexors and three extensors were targeted. The skin was first rubbed with alcohol. The electrodes were then coated with conductive gel and attached to the skin using double-sided tape. A single ground electrode was placed on the back of the neck.

### Data sampling & assembly

The data was assembled using Matlab R2013a. Kinematic and force data from the KINARM were sampled at 1 KHz. The recorded EMG signals were band-pass filtered online through the EMG system (20-450Hz) and sampled at 2 kHz. The EMG data was also high-pass filtered with a fifth-order, zero phase-lag Butterworth filter with a 30 Hz cutoff and then rectified. With regard to EMG, only data from probe trials were analyzed in the current study. To be able to compare and combine EMG data across muscles and subjects, each subject’s EMG data were normalized (z-transformed), similar to the procedure described elsewhere (Dimitriou, 2014; Dimitriou, Franklin, & Wolpert, 2012). Briefly, for each subject and muscle, all EMG signals pertaining to probe trials (N=128) were concatenated, and a grand mean and standard deviation was generated. These two numbers were then used to produce the normalized raw EMG data for each muscle of the subject (i.e., subtracting the grand mean and then dividing by the standard deviation). In addition to presenting and analyzing such data from individual muscles (e.g., Figure 3, left column), normalized EMG data across the recorded flexors (brachioradialis, biceps, posterior deltoid) and across the recorded extensors (short and long triceps, anterior deltoid) were collapsed (averaged) separately for each muscle group and subject. This produced one unified signal of ‘flexor’ and one for ‘extensor’ activity for each subject, which could then also be directly contrasted with the recorded endpoint forces applied on the robotic handle. To study stretch reflex responses to perturbations, the analyses focused on established time-periods, known to reflect spinal (monosynaptic) and transcortical stretch reflex loops (e.g., Hammond, 1956; Pruszynski et al., 2009; Scott, 2012; Scott et al., 2015). Specifically, using the onset of the kinematic perturbation to signify time zero, these periods were defined as the short-latency spinal ‘R1’ response (20 – 45 ms), and the long-latency ‘R2’ (50 – 75 ms) and ‘R3’ response (75 – 100 ms). A 50 ms interval (-50 – 0 ms) was chosen to represent pre-perturbation muscle activity (‘R0’), as defined elsewhere (e.g., Cluff & Scott, 2013). Voluntary EMG responses were considered to occur >120 ms following the onset of the kinematic perturbation (e.g., Pruszynski, Kurtzer, & Scott, 2008).

The force sensor embedded in the KINARM handle produced a signal reflecting the force applied by the hand in the principle axis of action during the postural perturbations (i.e., force along the Y axis), as well as a signal representing force applied in the X axis. Although stronger forces were expected along the Y axis, the imposed perturbations at the hand were expected to provoke reactive forces along the X axis as well (given the involuntary reflex action of the stretched muscles, coupled with the subjects’ initial posture; Figure 1A). Therefore both X and Y force channels were examined in the current study. The relevant analyses concentrated on a time period thought to correspond only to rapid (reflexively-produced) forces. Specifically, a minimum electromechanical delay of 30 ms is assumed between muscle electrical activity and actual force production (Ito, Murano, & Gomi, 2004). That is, the force equivalent of the ‘R1’, ‘R2’ and ‘R3’ reflex EMG responses are taken to be the “RF1” period (75-100 msec), “RF2” (100-125 msec) and “RF3” (125-150 msec), respectively. Accordingly, “RF0” was defined as force occurring in the period 0 - 30 msec. Direct contrast of reflexive EMG and force data indicate generally matching overall patterns in the current study (see e.g., Figure 8A vs. 8B). Any differences in statistical results obtained using force vs. EMG data are can be due to well-known differing levels of sensitivity and background noise in these recorded channels, or simply that actively-produced endpoint forces represent all involved antagonistic muscles.

The examination of probe/perturbation trials focused on analyzing EMG and force signals. However, in order to associate such data with the subject’s performance on the main learning task, the Initial Direction Error (IDE) was calculated using kinematic data from each voluntary movement (non-probe trials). As commonly defined, IDE represents the angular difference between the direction the hand was moving (calculated at initial peak velocity in each movement) and the direction subjects should have been moving to, given the target location. An IDE value was therefore produced for each trial of the main visuomotor learning task (i.e., non-probe trials). A common approach to examine kinematic learning progression involves fitting an exponential curve to the sequence of the generated IDEs to extract the error decay constant (‘learning rate’). In the current study, the calculated IDEs of each subject (N=384) were aligned and averaged across subjects, so that one mean IDE sequence was produced for the entirety of the task, separately for each experiment (Figure 2). Exponential curves were fitted separately to parts of the above sequence which corresponded to different task stages of each repetition (N=4 repetitions). In order to examine individual differences in the relationship between learning rate and reflex behavior, the calculated IDE’s for each movement were also used to obtain a single value representing learning rate (error decay constant) for each subject at ‘early exposure’ and ‘washout’. That is, separately for each of these two stages, the IDEs of a single subject were aligned and then averaged (i.e., mean across four IDE sequences for each stage, corresponding to the N=4 repetitions of the task). The above produced two independent sequences of IDE’s (each N=24 data-points) for each subject. Exponential fit on each sequence then produced a single learning rate for each subject for ‘early exposure’ and ‘washout’. The calculated IDEs and associated learning rates were the only pieces of information from non-probe trials that were used for statistical analyses.

### Statistical Analyses

Overall, most statistical analyses in the current study were performed on data that have had the equivalent ‘baseline’ state values subtracted, at the level of single subjects. This approach acted to both simplify the analyses and neutralize known main effects of experimental manipulations that have been previously applied outside the context of visuomotor learning (e.g., Marsden et al., 1976; Pruszynski et al., 2009; Pruszynski et al., 2008). Note, however, that plots of non-subtracted data (including ‘baseline’) are also presented throughout and the aforementioned effects are clearly reconstructed in the displayed data (e.g., see Figure 3). For statistical analyses involving EMG or force signals alone, only data across the well-known reflex/involuntary response periods were used (as described in the previous section). The relevant data used for each subject were averages across repetitions of a relevant trial type and across the time period in question. That is, a single data-point per subject, task stage, trial type and time period was ultimately generated for analysis purposes. The same general 3 × 2 × 4 repeated-measures ANOVA design was employed in order to analyze stretch reflex responses (as one would expect, these analyses only involved EMG data from stretching muscles). Specifically, in both experiments, a main factor in the repeated-measures ANOVA was task state, which had 3 levels: ‘early exposure’ vs. ‘late exposure’ vs. ‘washout’ (these values have had the equivalent ‘baseline’ value subtracted at the level of single subjects). Another main factor was the binary direction of the applied load and/or cued target (Figure 1DE), and the last factor was time period (4 levels: ‘R0’, ‘R1’, ‘R2’, ‘R3’ for EMG; or ‘RF0’, ‘RF1’, ‘RF2’, ‘RF3’ for force). If a significant main effect was found, Tukey’s HSD (honest significant difference) post hoc test was used, which takes into account multiple paired comparisons. As mentioned above, the force and EMG signals used in the ANOVA tests have had the equivalent ‘baseline’ values subtracted at the level of single subjects. Many of the results are therefore plotted relative to this nullifying baseline (e.g., Figure 4AB). Additional single-sample t-tests determined whether reflex responses at the subsequent stages of the visuomotor task significantly differed from those observed during ‘baseline’ (i.e., tested for significant difference from ‘0’). Statistical significance was considered at the p<0.05 level for all statistical tests. Correlations were also performed to examine relationships between individual learning rates (non-probe trials) and the magnitude of reflex EMG responses (probe trials).

**Figure 4.**
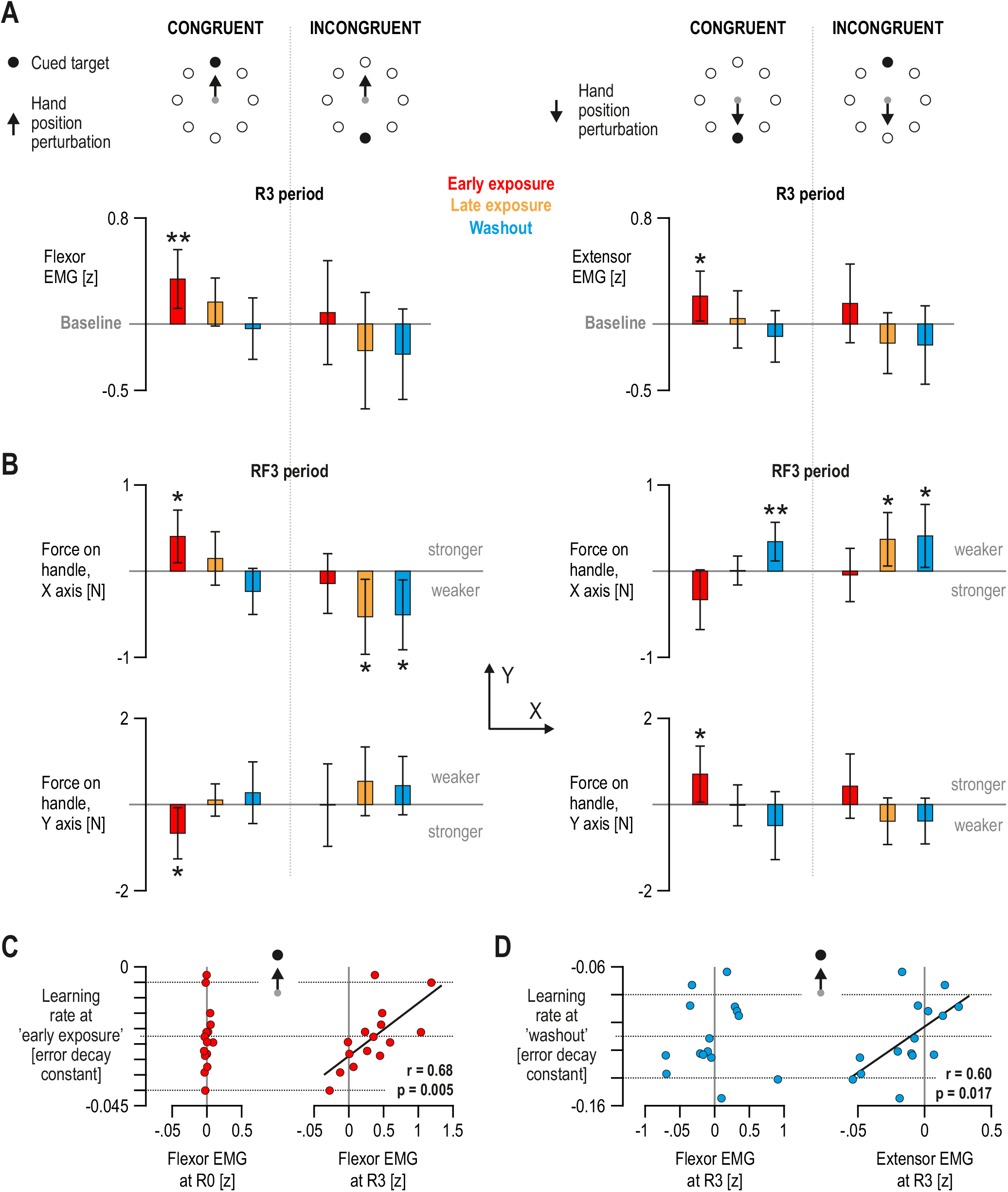
Modulation of long-latency reflex responses in Experiment 1. (**A**) Mean flexor EMG (left panel) and mean extensor EMG across all subjects (right panel), observed during the R3 stretch reflex period at the different stages of the main task, in response to congruent and incongruent haptic perturbations (see schematics). Throughout, error bars represent 95% confidence intervals. For both flexors and extensors, single-sample t-tests indicated significant deviation from ‘baseline’ responses only during the ‘early exposure’ stage, in response to ‘congruent’ perturbations. (**B**) As ‘A’, but referring to the reflexively-produced force (‘RF3’ period) generated by the subject’s hand on the robotic handle during the associated haptic perturbation. Single-sample t-tests complimented the results in ‘A’, but also indicated significantly weaker responses to the same congruent perturbation (extensor stretch) in the ‘washout’ stage (blue, right upper panel). ‘Stronger’ refers to a stronger response relative to the baseline state **(C, D)** Here, each dot represents a single subject. Only for cases when the hand was perturbed away from the torso, there is a significant relationship between long-latency responses of the stretching flexors (‘C’) and shortening extensors (‘D’) with individual learning rates, in ‘early exposure’ and ‘washout’, respectively (**= p<0.01, *= p<0.05).

## Results

The progression of movement errors in the visuomotor learning task (i.e., non-probe trials) of both Experiment 1 and 2 shows that subjects produced stereotypical adaptation behaviors (Figure 2). That is, exponential decay of movement error (i.e., learning) was evident in ‘early exposure’ (red) and ‘washout’ states (blue). The analysis of probe trials focused on time-periods known to reflect spinal and transcortical stretch reflex output (e.g., Hammond, 1956; Pruszynski et al., 2009; Scott, 2012; Scott et al., 2015). Visual inspection of single subject and across-subject signals in response to hand perturbations in Experiment 1 suggests that R3 EMG responses in the ‘early exposure’ state were higher than those observed in either ‘baseline’ or ‘washout’ (Figure 3). To statistically assess the above, a 3 (task state) × 2 (cued target direction) × 4 (time period: R0, R1, R2, R3) ANOVA was performed using flexor EMG. There was no main effect of cued target or time period (p>0.05). There was a significant main effect of task state on the magnitude of the flexor EMG responses (F(2,28)=4.63, p=0.018, η_p_^2^=0.25) and a significant interaction effect between time period and task state (F(6,84)=4.49, p=0.0006, η_p_^2^=0.24). Tukey’s post hoc test indicated that flexor EMG in the R3 period of ‘early exposure’ was significantly larger in magnitude than all other periods, including the corresponding R3 periods of ‘late exposure’ and ‘washout’, with p=0.0021 and p=0.00012, respectively. In addition, single-sample t-tests using flexor EMG at R3 revealed significantly larger values than the ‘baseline’ stage (i.e., >0; see Methods) only for perturbations applied at the ‘early exposure’ stage when the visual target direction was congruent (Figure 4A, left panel), with t(14)=3.29, p=0.0054.

Equivalent results were obtained for EMG from the stretching extensor muscles (Figure 4A, right panel). Complimentary results were also obtained when single sample t-tests examined the ‘long-latency’ reflexive forces exerted by the subjects on the robotic handle as a result of the ‘R3’ EMG response (i.e., forces at the ‘RF3’ interval: 120 -145 ms following perturbation onset). In addition to significantly larger forces -than baseline-in the ‘early exposure’ stage (Figure 4B), there were also significantly lower reflexive forces in the ‘washout’ state (‘X’ axis) for ‘congruent’ perturbations towards the torso (Figure 4B, right panel), with t(14)=3.28, p=0.006. Overall, consistent patterns emerged across flexor and extensor muscles, representing an upregulation of long-latency responses in ‘early exposure’ and an inhibition in the ‘washout’ state (Figure 4B). As Figure 4C shows, a positive relationship was found between individual learning rates at ‘early exposure’ and R3 flexor EMG, but no relationship between learning rate and pre-perturbation EMG levels (‘R0’). No equivalent relationship was found for extensor EMG. Interestingly, there was a significant relationship between individual learning rates in ‘washout’ and shortening extensor EMG at R3 (Figure 4D).

Figure 5 shows the brachioradialis EMG responses of two subjects when the hand was perturbed in Experiment 2: despite the direction of the imposed constant load and the location of the cued target, there are clear differences in monosynaptic EMG responses to the haptic perturbation as a function of visuomotor adaptation state. Most striking is the inhibition of monosynaptic R1 responses in the ‘washout’ state relative to ‘baseline’. Similar plots across all subjects revealed that this inhibition was clearly present in ~50% of subjects (the individual variability is addressed and accounted for below; i.e., Figure 7C). In addition, most subjects appeared to exhibit a consistent inhibition of their R3 response during ‘washout’ (Figure 6). An ANOVA test of the same design as in Experiment 1 (i.e., 3 × 2 × 4) indicated a main effect of task state on the responses of the stretching brachioradialis (F(2,28)=8.17, p=0.0016, η_p_^2^=0.37). Tukey’s test showed that responses during ‘early exposure’ were significantly larger than those in ‘late exposure’ and ‘washout’, with p=0.01 and p=0.0023, respectively. There was also an interaction effect between time period and task state (F(6,84)=2.84, p=0.014, η_p_^2^=0.19). Post hoc analysis revealed no significant differences in pre-perturbation (‘R0’) brachioradialis EMG across the task stages, but R3 responses in ‘late exposure’ and ‘washout’ were relatively smaller than all other cases (all p<0.002). In addition, the R2 response during ‘early exposure’ was significantly higher than that the R2 response during ‘washout’ (p=0.004). Single-sample t-tests produced complimentary findings (Figure 7A). In addition to significantly lower R3 EMG in ‘late exposure’ and ‘washout’, the monosynaptic R1 responses of the brachioradialis were also significantly smaller in ‘washout’ than baseline, regardless of the direction of the imposed load and cued target (p=0.043 and p=0.042: left vs. right panel; Figure 7A). Only the ‘congruent’ R2 responses during ‘early’ exposure’ significantly differed from baseline (p=0.029).

**Figure 5.**
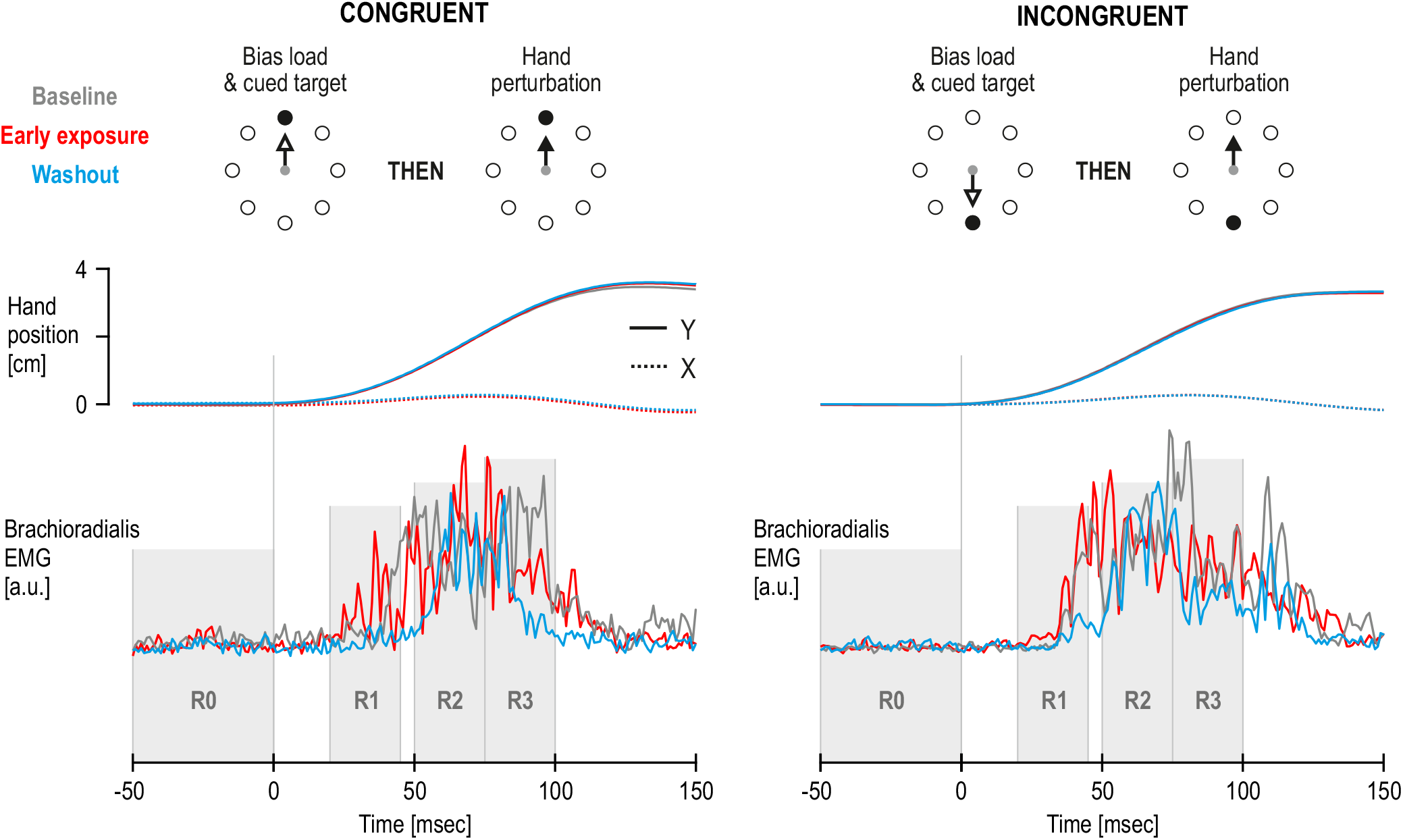
Single subject responses to hand perturbations in Experiment 2. Averaged responses across repetitions from two separate subjects (left vs. right column) when the hand was perturbed in the direction of flexor stretch, and the cued target location was either congruent (left column) or incongruent (right column; see schematics). Time ‘0’ represents the onset of the imposed kinematic perturbation. Despite virtually identical kinematics overall and same preperturbation EMG levels across task stages (‘R0’ period), there are clear differences in reflex EMG responses of both subjects as a function of task stage. This is true even for responses attributed to monosynaptic (spinal) reflex circuits (‘R1’). The most striking difference is the apparent inhibition of the R1 response during the ‘washout’ stage relative to ‘baseline’ (blue vs. gray). The above effect appeared despite that the task demanded of subjects to resist all perturbations equally across all task stages (i.e., subjects were instructed to remain immobile at the start point regardless, until the ‘Go’ visual signal was shown).

As mentioned above, the inhibition in R1 responses during ‘washout’ (e.g., Figure 5) was clearly present in ~50% of subjects; i.e., see ‘X’ axes in Figure 7B, second panel from left. However, across all subjects, I found a strong negative relationship between individual learning rates during ‘washout’ and corresponding R1 responses from the brachioradialis (Figure 7C, left), with r=-0.75 and p=0.0013. The same relationship was found for the aggregated ‘flexor’ EMG signal, with r=-0.65 and p=0.009 (Figure 7B, rightmost panel). That is, learning rates could account for the reflex behavior across all subjects, those that exhibited reflex inhibition and those that did not. Correlating the same learning rates with pre-perturbation activity (R0) of either the brachioradialis or across flexors indicated no relationship (Figure 7B). There was also no significant relationship between the same learning rates in ‘washout’ and corresponding flexor EMG responses in R2 (r=-0.28, p=0.31) or R3 periods (r=0.17, p=0.54).

**Figure 6.**
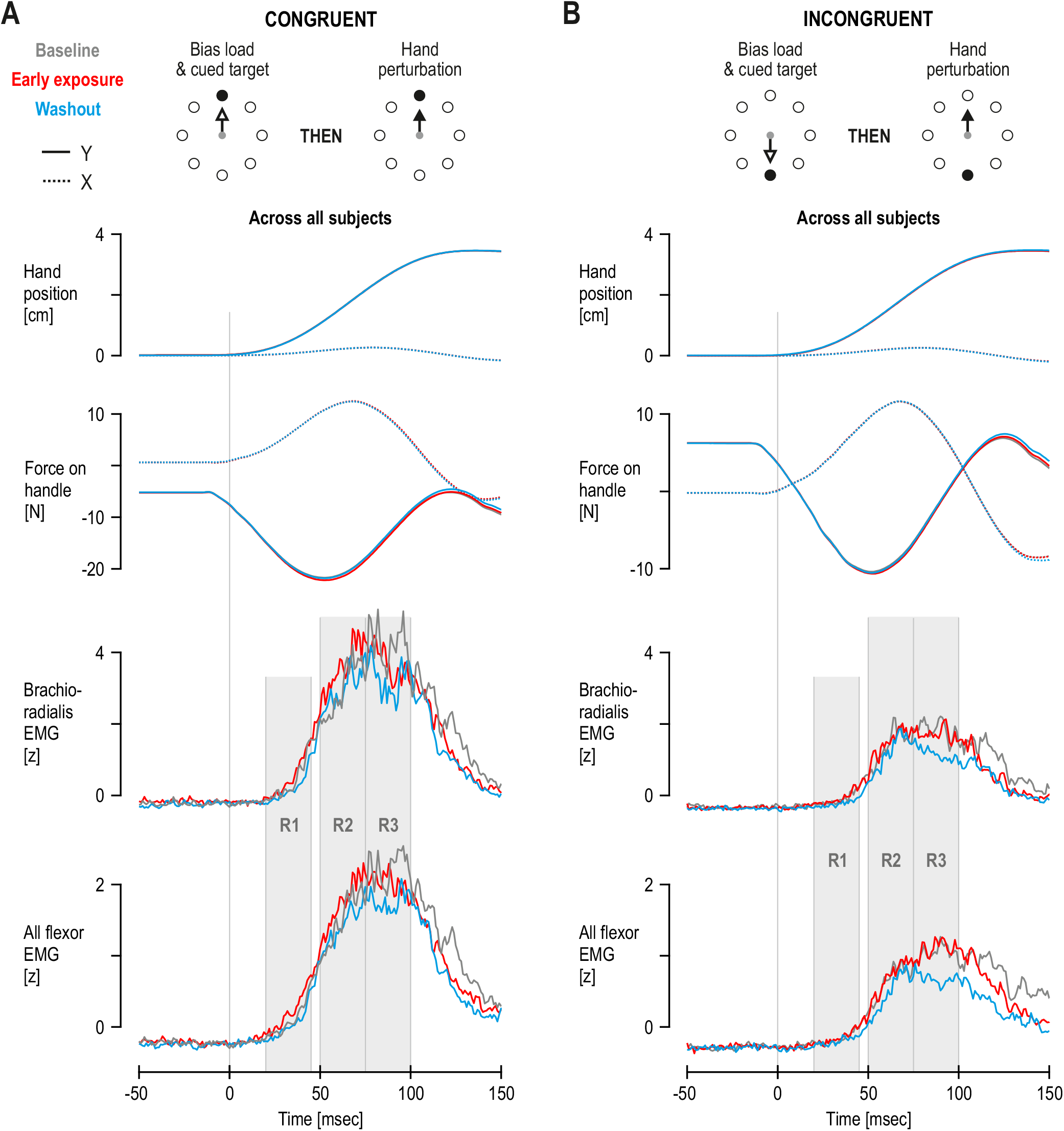
Responses to hand perturbations in Experiment 2. (A, B) As Figure 5, but here the traces represent means across all subjects (N=15). Time ‘0’ represents the onset of the imposed perturbation. Overall, the plots suggest that inhibition of the R1 response during ‘washout’ was strong in some subjects (e.g., Figure 5) but not others (i.e., ‘X’ axes in Figure 7C indicate that ~50% of all subjects exhibited inhibition of R1; the variability across subjects is also addressed in Figure 7C). A much more consistent inhibition of the R3 response was evident across subjects during ‘washout’, particularly in response to incongruent perturbations (‘B’). Error bars are omitted for visual clarity but variances across subjects are tested statistically and also displayed in Figure 7.

**Figure 7.**
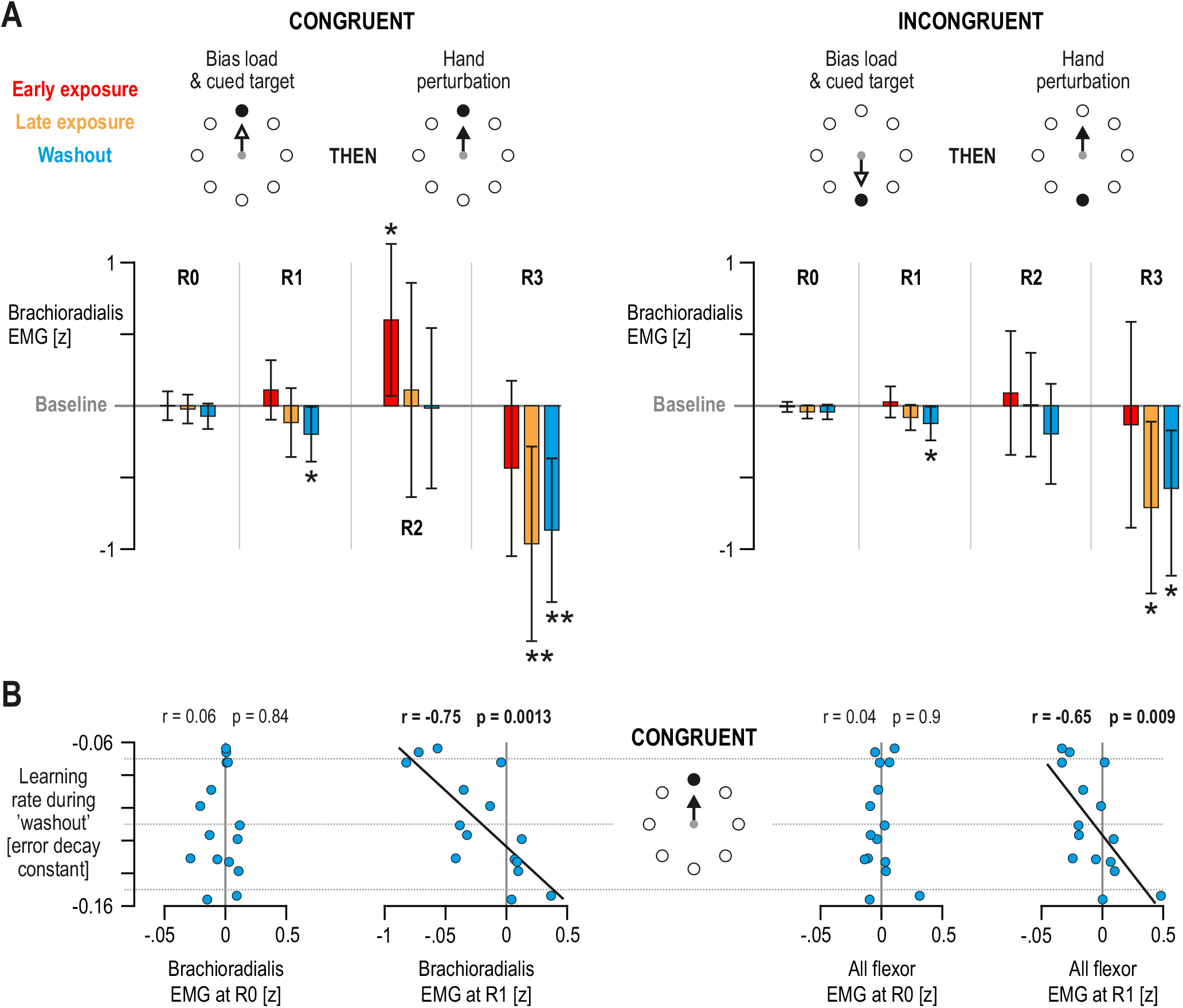
Task-dependent modulation of short- and long-latency stretch reflexes of distal muscles (**A**) Mean brachioradialis EMG across subjects at the different reflex latencies, in response to congruent (left panel) or incongruent perturbations of the hand (right panel). The perturbations were applied in the direction of flexor stretch (see schematics). Averaged preperturbation values (‘R0’) are also shown. Error bars represent 95% confidence intervals. As in Figure 4, the data is presented with reference to the individual ‘baseline’ responses (i.e., individual responses at the ‘baseline’ stage of the visuomotor task). Overall, single-sample t-tests indicated significant decrease in R3 magnitude both in the ‘washout’ (blue) and ‘late exposure’ periods (orange), but only in the ‘washout’ phase was there a significant inhibition in monosynaptic R1 responses. (**B**) Here, each dot represents data from a single subject (N=15). There was a significant relationship between individual learning rates in ‘washout’ and brachioradialis EMG at R1, observed in response to congruent perturbations in this stage. There was no significant relationship between the same learning rates and EMG levels at R0 (leftmost panel). Note that the subjects showing the greater inhibition in R1 (and worst learning performance) were not the ones with weaker responses than baseline in R0. This further validates the inhibitory R1 effect in ‘A’ (top left panel) as a task-dependent phenomenon. Equivalent relationships were found when the aggregated flexor EMG signal was used instead of the brachioradialis (right panels). There were no significant relationships between these learning rates and R2 or R3 responses of the brachioradialis or the aggregated flexor signal (**= p<0.01, *= p<0.05).

Similar response patterns were observed from the aggregated ‘flexor’ EMG signal when the same ANOVA test as above (i.e., 3 × 2 × 4) was performed. Specifically, there was main effect of task state on reflexive ‘flexor’ EMG responses (F(2,28)=8.68, p=0.0011, η_p_^2^=0.38). Tukey’s test showed that responses during ‘early exposure’ were significantly larger than those in ‘late exposure’ and ‘washout’, with p=0.006 and p=0.002, respectively. There was also an interaction effect between time period and task state (F(6,84)=4.37, p=0.0007, η_p_^2^=0.24). Post hoc analysis revealed no significant differences in pre-perturbation (‘R0’) ‘flexor ‘EMG across the task stages, but R3 responses in ‘late exposure’ and ‘washout’ were relatively smaller than all other cases (all p<0.0001). Single-sample t-tests produced largely complimentary findings (Figure 8A), such as significantly lower R3 EMG than ‘baseline’ (i.e., than ‘0’) in ‘late exposure’ and ‘washout’. The decrease in R1 response during ‘washout’ (as observed for the brachioradialis, Figure 7A) was not deemed significant in this case. Instead, a near-significant increase in R1 EMG responses was observed in ‘early exposure’ (p=0.06; Figure 8A, left panel). When the same test was applied using the equivalent reflexively-produced force along the ‘X’ axis (‘RF1’), this deviation did reach statistical significance, with t(14)=2.18, p=0.045 (Figure 8B, left panel). Note that the majority of the recorded muscles were proximal ones (except brachioradialis), therefore the produced endpoint forces are expected to represent more closely the ‘aggregated’ EMG signals. There was also significantly smaller ‘RF3’ force in ‘washout’ compared to ‘baseline’, in response to ‘incongruent perturbations’, with t(14)=-2.37, p=0.033 (Figure 8B, right panel). Same ANOVA design as above, produced a significant interaction effect between time period and task state on reflexive force along the ‘X’ axis (F(6,84)=2.77, p=0.0165, η_p_^2^=0.17), and an interaction effect between period, task state and perturbation congruence (F(6,84)=2.57, p=0.024, η_p_^2^=0.16). Post hoc analysis revealed no significant differences in pre-perturbation force (‘RF0’) across the task stages, in either type of perturbation. As a contrast between the two panels of Figure 8 suggests, post hoc analyses indicated that relative increases of reflexive force (‘X’ axis) in ‘early exposure’ primarily involved responses to ‘congruent’ perturbations, and the largest decrease in ‘RF3’ was at the ‘washout’ state for ‘incongruent’ perturbations.

**Figure 8.**
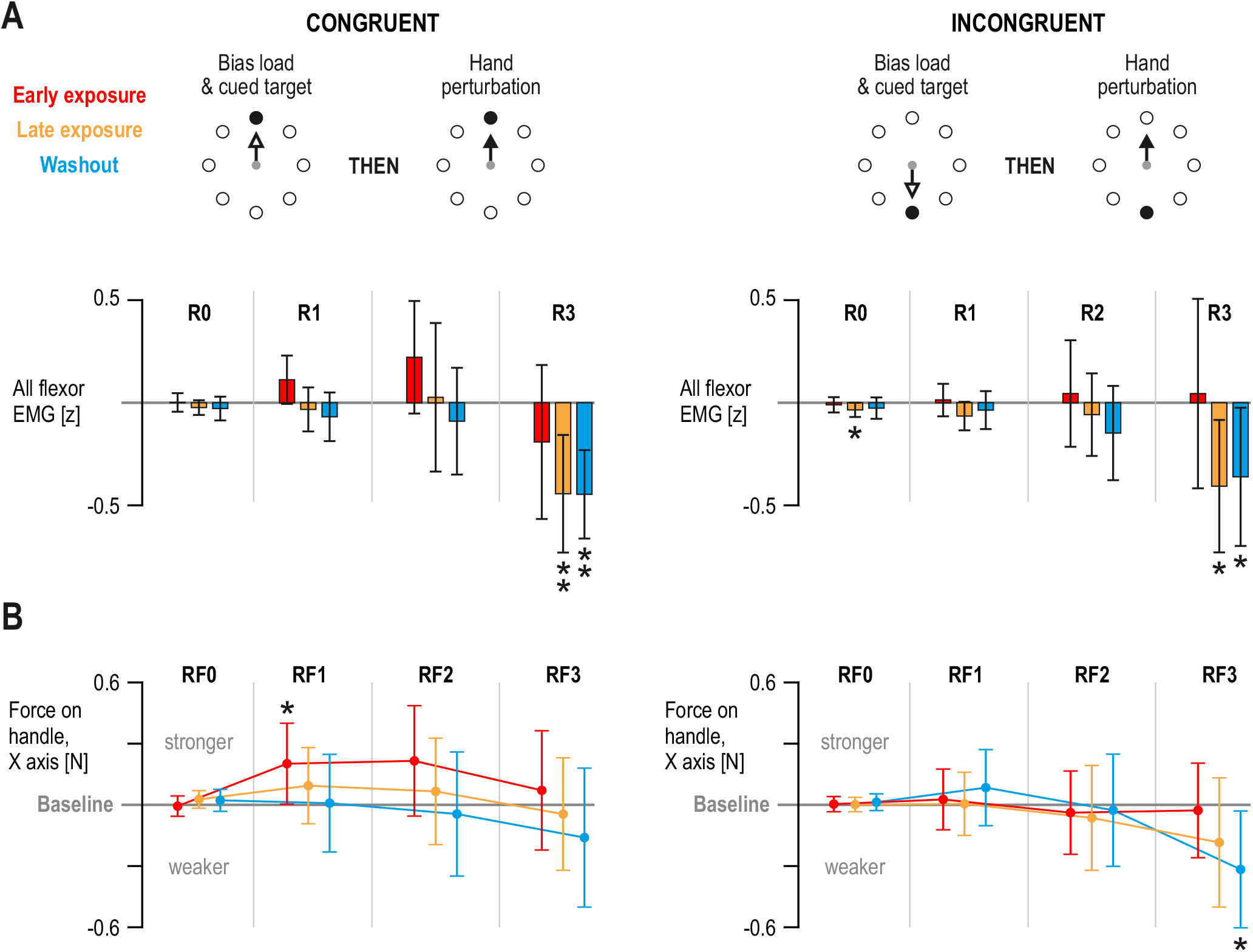
*Task-dependent modulation of short- and long-latency stretch reflexes* (A) As Figure 7A, but here representing the aggregated (average) of normalized EMG signals across all recorded flexor muscles (majority were proximal muscles). The results indicate a relative increase in R1 of proximal muscles in the ’early exposure stage’, but statistical significance was reached only when the equivalent endpoint forces were examined (see ‘B’) (B) The end-point force equivalent of ’A’ (’X’ axis). (**= p<0.01, *= p<0.05).

An ANOVA test using force produced along the ‘Y’ axis as a result of perturbations away from the torso, revealed no significant effects relating to task state. However, an equivalent analysis of ‘Y’ force produced as a result of perturbations towards the torso did reveal some clear differences (Figure 9). Specifically, ANOVA showed a main effect of task state on force along the ‘Y’ axis, with F(2,28)=3.8, p=0.034, η_p_^2^=0.21. Tukey’s test showed that responses during ‘early exposure’ were significantly larger than those in ‘washout’, with p=0.04. There was also an interaction effect between time period and task state (F(6,84)=2.74, p=0.017, η_p_^2^=0.16). Post hoc analysis revealed no significant differences in ‘RF0’ force across the task stages, but most significant differences involved a relative decrease of reflexively-produced forces during ‘washout’. Single-sample t-tests produced complimentary findings. As Figure 9 shows, a consistent effect involves a decrease in responses during the ‘washout’ state, but reaching significance only in the ‘long-latency’ periods ‘RF2’ and ‘RF3’.

**Figure 9.**
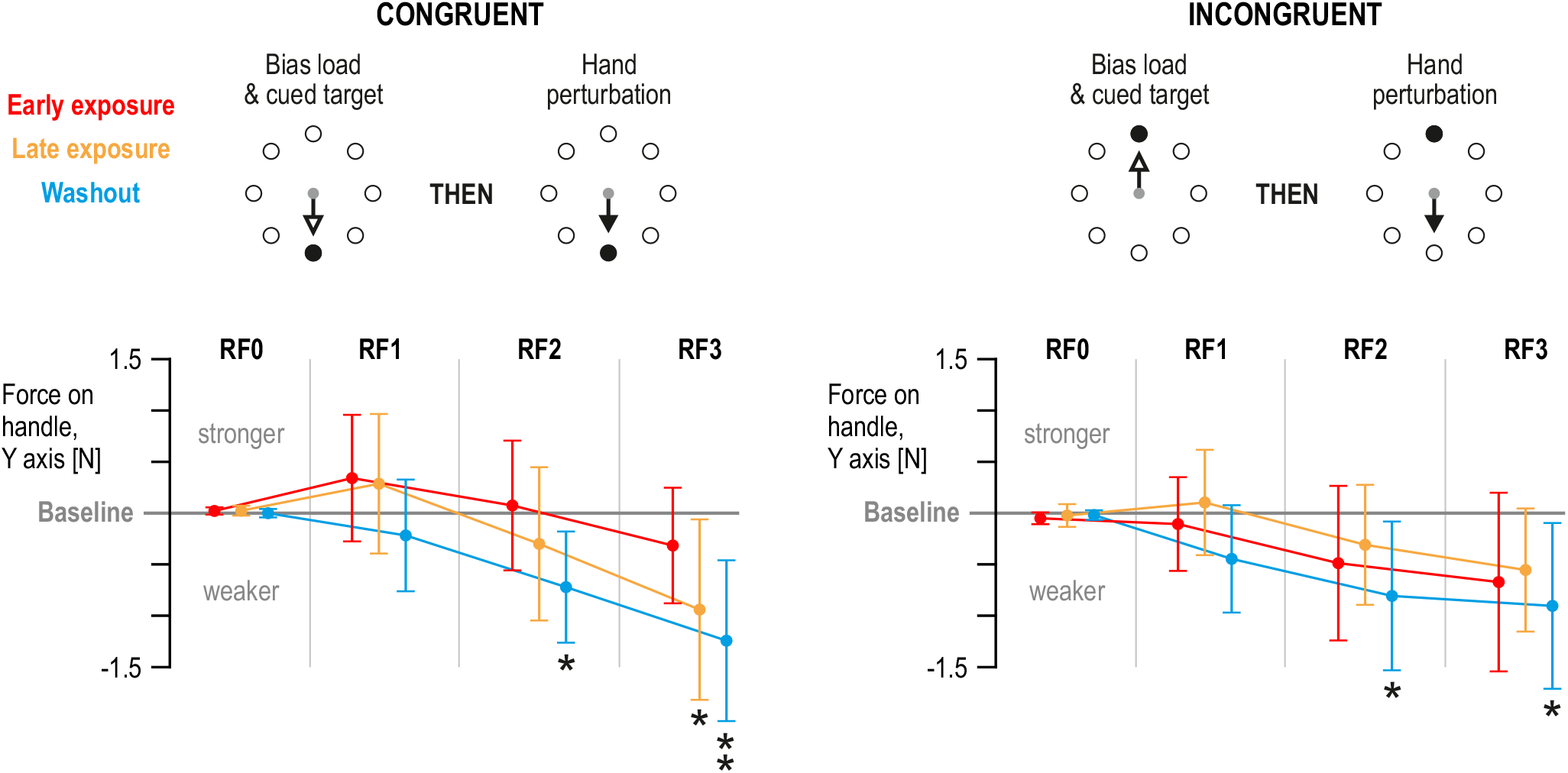
Task-dependent modulation of feedback responses induced by perturbations towards the torso (A) As Figure 8B, but here representing the end-point force responses (’Y axis’) to haptic perturbations in the direction of extensor stretch. A similar tendency of increased RF1 forces is observed in the ’congruent’ case (left), as in Figure 8B, but here statistical significance was not reached. However, consistent and significant decreases of ‘long-latency’ reflexive forces are observed in the ‘washout’ state, regardless of perturbation congruence and immediate task goal i.e., to resist the perturbation (**= p<0.01, *= p<0.05).

## Discussion

The current study examined the modulation of stretch reflex responses to position perturbations of the hand, given recent evidence of task-dependency in proprioceptive afferent signals during movement in visuomotor learning (Dimitriou, 2016). Here, subjects performed a classic visuomotor rotation task by making center-out reaching movements with their right hand. On randomly interleaved probe trials within the different stages of this learning task, the position of the hand was unpredictably perturbed during the movement preparation stage (i.e., while subjects were waiting for a ‘Go’ signal to move). In addition to any upregulation of stretch reflex responses, it was expected that feedback gains should also exhibit a task-dependent inhibition at certain stages, independent of the muscle’s contractile state pre-perturbation. Such behavior was expected to occur at monosynaptic (‘R1’) latencies as well. The current study also allowed for an examination of individual differences in the relationship between reflex gain modulation (probe trials) and motor learning performance (non-probe trials). To summarize, it appears that motor learning can indeed involve inhibition of reflex responses, and the monosynaptic reflex output (‘R1’) can also be affected by higher-level aspects of a sensorimotor task (e.g., Figures 5–8). For more distal muscles (brachioradialis), a close relationship was also found between R1 reflex gains and individual learning rates. All haptic perturbations in the current study were applied when the subjects were completely immobile and waiting for the ‘Go’ signal in order to initiate movement. Therefore, as evidenced by the recorded EMG and kinematics in the pre-perturbation period (‘R0’; e.g., Figures 3, 5 & 6), the position of the hand was perturbed before any transition to movement was either cued, generated or even desired. It is known that when faced with novel dynamics, the nervous system ‘loads’ task-relevant feedback controllers already at the movement preparation stage (Ahmadi-Pajouh et al., 2012). The current study indicates that an equivalent ‘pre-loading’ of feedback controllers can also occur in kinematic learning.

Overall, the demonstrated inhibition in reflex responses as a function of adaptation state - including at long-latency intervals-reflects a higher degree of sophistication in feedback responses than previously thought. That is, being in a state of motor adaptation (i.e., learning) has been previously shown to involve an upregulation of the transcortical component of rapid feedback responses (e.g., Ahmadi-Pajouh et al., 2012; Cluff & Scott, 2013; Franklin et al., 2012). It has been suggested that uncertainty in the state of the body, such as that arising through interaction with a novel environment, leads to an upregulation of feedback gains which in turn allow the system to minimize movement error or disturbances while the feedforward controller adapts to the new state of affairs. This kind of upregulation in feedback gains was also found in the current study, when subjects were initially exposed to the visual distortion i.e., during the ‘early exposure’ stage (e.g., Figures 4, 7 & 8). However, as mentioned above, a decrease in feedback gains was consistently found in the ‘washout’ stage. Simply reversing the assumption for the purpose of upregulation in feedback gains, leads one to assume that more disturbance or movement error (i.e., variability) is actively promoted by the system during the ‘washout’ state, relative to ‘baseline’. Indeed, recent evidence has revealed a positive role for task-relevant motor variability (i.e., ‘exploration’) in facilitation of motor learning in humans (Wu, Miyamoto, Gonzalez Castro, Olveczky, & Smith, 2014). If variability has a positive impact on learning performance, then one may wonder why the system prefers to upregulate gains in certain circumstances, such as during ‘early exposure’ to an altered kinematic environment (e.g., Figures 4 & 7). Possibly, the reason lies with the task-relevance of the sensory inflow itself. That is, in the ‘washout’ stage, the proprioceptive afferent feedback is directly reflective of the task-relevant consequences of the motor commands: proprioceptive information about movement direction and position of the hand is congruent with the direction and position of the visual cursor. More ‘exploration’ by the limb in this case may allow for faster update of internal models based on proprioceptive information. The situation is likely more complicated in dynamic motor learning, where the altered environment itself can cause proprioceptive signals to change. Irrespective of the direction of feedback modulation in kinematic ‘washout’, the documented modulation of reflex gains in this stage (relative to ‘baseline’) also adds to existing claims that the kinematic “unlearning” process is actually an active process, involving more than just passive memory decay (Dimitriou, 2016; Kitago, Ryan, Mazzoni, Krakauer, & Haith, 2013).

At least with respect to maintenance of body configuration (‘posture’), it is has been generally believed that task-dependent control is achieved by adjusting voluntary motor activity and transcortical (‘long-latency’) reflexes, and that spinal monosynaptic circuits are not engaged in such flexible task-level control (e.g., Hammond, 1956; Marsden et al., 1976; Pruszynski et al., 2009; Pruszynski & Scott, 2012; Scott, 2012; Scott et al., 2015). To my knowledge, the current results represent the first systematic demonstration of task-dependent modulation in human postural monosynaptic reflexes (e.g., Figures 5, 7 & 8). Such effects appeared systematically only in Experiment 2. The detection of segmental effects in Experiment 2 vs. Experiment 1 was most probably facilitated by the pre-loading of muscles, a manipulation which is known to help detection of short-latency EMG signals (Marsden et al., 1976; Pruszynski et al., 2009). Indeed, ‘R1’ responses were generally weak or absent in all task stages in Experiment 1 (see Figure 3). In Experiment 2, pre-loading of muscles brought about an expected overall increase in gain (i.e., R1 responses appeared consistently across task stages, including ‘baseline’; e.g., Figure 5–6), which in turn allowed differences in R1 to be detected across the different visuomotor task stages. Even though task-dependent differences occurred only when the subjects also faced an external mechanical load, the perturbed muscle’s state was kept the same throughout all probe trials, in order to isolate any effect of task state/stage on reflex gains. In other words, the task-dependent effects across task stages were not due to the state of the muscle itself. An inhibition of R1 responses was only found in more distal muscles (brachioradialis), whereas more proximal muscles represented a relative increase in R1. This difference likely reflects alternate descending innervation. That is, it is well known that distal (e.g., forearm) skeletal muscles of the primate are also innervated with more direct connections from the cortex. This includes monosynaptic excitatory and di-synaptic inhibitory connections upon fusimotor neurons controlling muscle spindle output (Clough, Phillips, & Sheridan, 1971). This may have affected the R1 output of distal muscles, which was also the one more closely related to individual learning rates in the visuomotor task (Figure 7B).

Interestingly, motor learning performance (i.e., adaptation rate) in Experiment 2 could account for the modulation of R1 feedback gains across all subjects: those which displayed a modulation (inhibition) of the monosynaptic reflex and those that did not (r=-0.75, p=0.0013; Figure 7B). It is therefore not surprising that the overall inhibitory effect of the ‘washout’ state on reflex EMG output was marginally statistically significant across all subjects. Inspecting the R1 responses of each subject during ‘washout’ (i.e., X axes in Figure 7B, second panel from the left), reveals a substantial decrease in monosynaptic reflex gain for the subjects displaying relatively worse motor learning performance. This does not necessarily imply that weaker segmental gains lead to worse motor learning performance. As mentioned above, weaker reflexes can be beneficial (Wu et al., 2014) allowing for more ‘exploration’ along task-relevant dimensions. The inhibition of R1 gains in the current study may possibly reflect a compensatory process such as proprioceptor tuning through independent fusimotor control (Dimitriou, 2016), helping to maintain a certain level of performance. Indeed, as normally the case, learning performance in this study was universally better in ‘washout’ than the ‘early exposure’ stage (e.g., Figure 4C vs. 4D). Decreasing the muscle spindle’s sensitivity to the first and/or second derivative of muscle length may have led to the observed modulation in the spinal feedback gains. This may have involved a task-dependent decrease in ‘dynamic’ fusimotor output and/or an increase in static fusimotor activity. However, other mechanisms such as state-dependent neuromodulation (e.g., Marder, O’Leary, & Shruti, 2014) cannot be excluded. But such neuromodulation would probably lead to a universal decrease (or increase) in reflex feedback gains, whereas the current results reveal a non-uniform modulation. For example, Figure 7A (left) indicates a significant decrease in ‘R1’ and ‘R3’ responses in ‘washout’ compared to ‘baseline’, whereas no consistent deviation is observed in the ‘R2’ period. Fusimotor control may allow for relatively more flexibility in rapid feedback responses. For example, it is possible that feedback mechanisms associated with ‘R3’ responses are more heavily dependent on primary (‘Ia’) muscle spindle output than the ‘R2’ response.

A link between segmental reflex modulation and individual motor learning performance occurred only when the former was measured in ‘congruent’ probe trials: perturbations whose direction was congruent with the direction of the visually cued target. No relationship was found when the same learning rates where correlated with reflex gains measured during ‘incongruent’ perturbations (i.e., the hand was perturbed in the direction opposite to the cued target). The presence of a relationship between segmental reflex gains and performance in congruent trials may simply reflect the ecologically validity of this particular kind of probe trial. In everyday life, we normally identify the location of a desired visual target and then reach towards. That is, the direction of the movement is congruent with the direction of the identified target. Our nervous system may purposefully and habitually allow segmental feedback gains to modulate according to learning rate only along the goal-relevant direction. Indeed, both the current study (e.g., Figure 3B) and elsewhere (Pruszynski et al., 2008), perturbing the limb in a direction away from the highlighted target leads to a universal increase in feedback gains, a process that may largely override any effects of motor adaptation rate. However, once this general effect was nullified by subtracting the response magnitude observed at the ‘baseline’ state, a further effect was revealed: incongruence lead to a relative decrease in feedback responses, with the biggest difference seen in the ‘late exposure’ stage (see e.g., Figure 4B, top left).

To summarize, the current results demonstrate that the system’s ‘control policy’ in motor adaptation can include a state-dependent inhibition of stretch reflexes, in addition to any upregulation of feedback gains. Moreover, the current study shows that aspects of this policy can affect the output of monosynaptic feedback circuits. For more distal muscles (brachioradialis), there was a task-dependent inhibition of the monosynaptic reflex response, and the R1 gains across all subjects reflected individual visuomotor learning performance. Task-dependent modulation of R1 responses points to a form of state-dependent decentralized control that can tune spinal circuits according to task-level dimensions. It has been generally assumed that segmental control of posture is unaffected by the system’s control policy, suggesting that compensation for this ‘unruly’ output might even be warranted. The current results lead to the conclusion that a certain degree of flexible tuning is possible at the spinal level. In other words, the system’s policy of how to perform a motor action can apparently extend to segmental circuits flexibly (i.e., within the timeframe of a single experimental session) and may well include a plan for independent and task-specific modulation of fusimotor neurons affecting muscle afferents.

## Acknowledgements

I would like to thank Kamil Antos for assistance with programming the robotic manipulandum, Carola Hjältén for technical assistance, as well as S. Papaioannou and D.W. Franklin for comments on the initial draft of the manuscript.

## References

Ahmadi-Pajouh, M. A., Towhidkhah, F., & Shadmehr, R. (2012). Preparing to reach: selecting an adaptive long-latency feedback controller. The Journal of Neuroscience, 32(28), 9537–9545.

Akazawa, K., Aldridge, J. W., Steeves, J. D., & Stein, R. B. (1982). Modulation of stretch reflexes during locomotion in the mesencephalic cat. Journal of Physiology, 329, 553–567.

Capaday, C., & Stein, R. B. (1986). Amplitude modulation of the soleus H-reflex in the human during walking and standing. The Journal of Neuroscience, 6(5), 1308–1313.

Clough, J. F., Phillips, C. G., & Sheridan, J. D. (1971). The short-latency projection from the baboon’s motor cortex to fusimotor neurones of the forearm and hand. Journal of Physiology, 216(2), 257–279.

Cluff, T., & Scott, S. H. (2013). Rapid feedback responses correlate with reach adaptation and properties of novel upper limb loads. The Journal of Neuroscience, 33(40), 15903–15914. doi:10.1523/JNEUROSCI.0263-13.2013

Dimitriou, M. (2014). Human Muscle Spindle Sensitivity Reflects the Balance of Activity between Antagonistic Muscles. The Journal of Neuroscience, 34(41), 13644–13655. doi:10.1523/JNEUROSCI.2611-14.2014

Dimitriou, M. (2016). Enhanced Muscle Afferent Signals during Motor Learning in Humans. Current Biology, 26(8), 1062–1068. doi:10.1016/j.cub.2016.02.030

Dimitriou, M., Franklin, D. W., & Wolpert, D. M. (2012). Task-dependent coordination of rapid bimanual motor responses. Journal of Neurophysiology, 107(3), 890–901.

Dimitriou, M., Wolpert, D. M., & Franklin, D. W. (2013). The temporal evolution of feedback gains rapidly update to task demands. The Journal of Neuroscience, 33(26), 10898–10909.

Franklin, S., Wolpert, D. M., & Franklin, D. W. (2012). Visuomotor feedback gains upregulate during the learning of novel dynamics. Journal of Neurophysiology, 108(2), 467–478.

Hammond, P. H. (1956). The influence of prior instruction to the subject on an apparently involuntary neuro-muscular response. Journal of Physiology, 132(1), 17–18P.

Howard, I. S., Ingram, J. N., & Wolpert, D. M. (2009). A modular planar robotic manipulandum with end-point torque control. Journal of Neuroscience Methods, 181(2), 199–211. doi:10.1016/j.jneumeth.2009.05.005

Ito, T., Murano, E. Z., & Gomi, H. (2004). Fast force-generation dynamics of human articulatory muscles. Journal of Applied Physiology (1985), 96(6), 2318–2324; discussion 2317. doi:10.1152/japplphysiol.01048.2003

Kitago, T., Ryan, S. L., Mazzoni, P., Krakauer, J. W., & Haith, A. M. (2013). Unlearning versus savings in visuomotor adaptation: comparing effects of washout, passage of time, and removal of errors on motor memory. Frontiers in Human Neuroscience, 7, 307. doi:10.3389/fnhum.2013.00307

Marder, E., O’Leary, T., & Shruti, S. (2014). Neuromodulation of circuits with variable parameters: single neurons and small circuits reveal principles of state-dependent and robust neuromodulation. Annual Review of Neuroscience, 37, 329–346. doi:10.1146/annurev-neuro-071013-013958

Marsden, C. D., Merton, P. A., & Morton, H. B. (1976). Servo action in the human thumb. Journal of Physiology, 257(1), 1–44.

Mortimer, J. A., Webster, D. D., & Dukich, T. G. (1981). Changes in short and long latency stretch responses during the transition from posture to movement. Brain Research, 229(2), 337–351.

Pruszynski, J. A., Kurtzer, I., Lillicrap, T. P., & Scott, S. H. (2009). Temporal evolution of “automatic gain-scaling”. Journal of Neurophysiology, 102(2), 992–1003.

Pruszynski, J. A., Kurtzer, I., & Scott, S. H. (2008). Rapid motor responses are appropriately tuned to the metrics of a visuospatial task. Journal of Neurophysiology, 100(1), 224–238. doi:10.1152/jn.90262.2008

Pruszynski, J. A., & Scott, S. H. (2012). Optimal feedback control and the long-latency stretch response. Experimental Brain Ressearch, 218(3), 341–359.

Scott, S. H. (2012). The computational and neural basis of voluntary motor control and planning. Trends in Cognitive Sciences, 16(11), 541–549.

Scott, S. H. (2016). A Functional Taxonomy of Bottom-Up Sensory Feedback Processing for Motor Actions. Trends in Neurosciences, 39(8), 512–526. doi:10.1016/j.tins.2016.06.001

Scott, S. H., Cluff, T., Lowrey, C. R., & Takei, T. (2015). Feedback control during voluntary motor actions. Current Opinion in Neurobiology, 33, 85–94. doi:10.1016/j.conb.2015.03.006

Wolpaw, J. R. (1982). Change in short-latency response to limb displacement in primates. Federation Proceedings, 41(6), 2156–2159.

Wu, H. G., Miyamoto, Y. R., Gonzalez Castro, L. N., Olveczky, B. P., & Smith, M. A. (2014). Temporal structure of motor variability is dynamically regulated and predicts motor learning ability. Nature Neuroscience, 17(2), 312–321. doi:10.1038/nn.3616

